# Voltage-dependent K^+^ Channels Improve the Energy Efficiency of Signalling in Blowfly Photoreceptors

**DOI:** 10.1101/090308

**Authors:** Francisco J. H. Heras, John Anderson, Simon B. Laughlin, Jeremy E. Niven

## Abstract

Voltage-dependent conductances in many spiking neurons are tuned to reduce action potential energy consumption, so improving the energy efficiency of spike coding. However, the contribution of voltage-dependent conductances to the energy efficiency of analogue coding, by graded potentials in dendrites and non-spiking neurons, remains unclear. We investigate the contribution of voltage-dependent conductances to the energy efficiency of analogue coding by modelling blowfly R1-6 photoreceptor membrane. Two voltage-dependent delayed rectifier K^+^ conductances (DRs) shape the membrane's voltage response and contribute to light adaptation. They make two types of energy saving. By reducing membrane resistance upon depolarisation they convert the cheap, low bandwidth membrane needed in dim light to the expensive high bandwidth membrane needed in bright light. This investment of energy in bandwidth according to functional requirements can halve daily energy consumption. Second, DRs produce negative feedback that reduces membrane impedance and increases bandwidth. This negative feedback allows an active membrane with DRs to consume at least 30% less energy than a passive membrane with the same capacitance and bandwidth. Voltage gated conductances in other non-spiking neurons, and in dendrites, might be organized to make similar savings.

## 1 Introduction

Energy efficiency is a fundamental organizational principle in brains that influences numerous aspects of signal generation, processing and transmission [1, 2, 3, 4, 5]. Most of the energy that supports electrical signalling is consumed by ion pumps to restore the large numbers of ions that move across the membrane. Consequently, energy efficiency can be substantially improved by adapting and organizing a neuron’s ionic conductances to reduce the flux required to produce a signal of necessary quality. To this end, a number of neurons configure voltage-dependent Na^+^ and K^+^ conductances to improve action potential energy efficiency by reducing the overlap between Na^+^ influx and K^+^ efflux [6, 7, 8, 3, 9]. With more efficient action potentials, circuits expend less energy on transmission and more energy on processing analogue synaptic signals [8, 10].

Voltage-dependent conductances also participate in the processing of analogue signals, both in the dendrites of spiking neurons (for a review see [11]), and in many types of non-spiking neuron, such as photoreceptors (e.g. [12, 13]), second-order visual interneurons (e.g. [14, 15]), auditory hair cells [16], and insect pre-motor interneurons [17]. An early study of fruit fly photoreceptors suggested that a voltage-dependent conductance could increase the energy efficiency of analogue processing [18]. Here we model the photoreceptor membrane of the blowfly to determine how and to what extent delayed rectifier voltage-dependent K^+^ conductances increase the energy efficiency of analogue processing.

Blowfly R1-6 photoreceptors employ a fast and a slow delayed rectifier conductance (FDR and SDR, respectively) to adapt the membrane gain and bandwidth to the background light level [13]. Increasing levels of background light progressively depolarise the photoreceptor from a dark resting potential of approximately −60 mV to approximately −30 mV in full daylight. The photoreceptor adapts from darkness to full daylight by progressively reducing the sensitivity of its voltage response and increasing its voltage signal bandwidth. The cut-off frequency of the voltage signal, the frequency at which the power spectrum falls to half-maximum, increases from 20 Hz in dark-adapted to 80 Hz in fully light-adapted photoreceptors [19]. This 4-fold increase is achieved primarily by accelerating the kinetics of the processes that link the activation of a rhodopsin molecule by a photon to the passage of light-induced current (LiC) through trp channels [20]. By progressively activating with increasing depolarisation, and hence light level, FDR and SDR assist light adaptation in several ways: (i) they oppose the depolarization produced by the LiC, to reduce voltage saturation [13]; (ii) they lower membrane impedance to reduce membrane gain (change in voltage signal amplitude/change in LiC), thereby reducing sensitivity [13]; (iii) they reduce the membrane time constant to expand the membrane bandwidth in step with the bandwidth of the light response [13]; (iv) the FDR produces shunt peaking, increasing the gain-bandwidth product of the membrane [21]; and (v) the SDR converts the membrane’s frequency response from low-pass at low light levels to band-pass by activating at high light levels. It has also been suggested that the progressive activation of FDR and SDR increases energy efficiency [22], but this has not been studied systematically. We use a Hodgkin-Huxley model of the blowfly photoreceptor membrane to explore the energy savings made by its voltage-dependent K^+^ conductances. We identify and evaluate two types of saving. First, voltage-dependency avoids excessive energy consumption by regulating K^+^ conductance according to functional requirements; a high conductance when depolarised by bright light to provide a low gain and high bandwidth, and a low conductance when close to the dark resting potential to provide a high gain and low bandwidth. We show that this voltage-dependent reduction in conductance from light to dark reduces energy consumption more than 6-fold. The second type of saving is less obvious, but nonetheless significant. It arises because voltage-dependent K^+^ conductances actively counteract the change in membrane potential produced by a change in LiC with a flow of current of opposite polarity. This negative feedback reduces the amplitude and time-to-peak of the voltage response, thereby increasing the membrane bandwidth. By comparing the energy consumed by currents crossing a membrane containing voltage-dependent K^+^ conductances to the consumption of membrane that achieves an identical bandwidth using a voltage-independent K^+^ conductance, we find that the active membrane consumes 50% less energy than the passive. Thus, voltage-dependent K^+^ conductances similar to those found in the majority of neurons increase the energy efficiency of analogue processing by reducing the cost of an expensive necessity, sufficient bandwidth.

## 2 Methods

### 2.1 Delayed rectifier characteristics

We used a Hodgkin-Huxley [23] (HH) type model of the two delayed rectifier conductances (DRs) present in blowfly, *Calliphora vicina*, R1-6 photoreceptors. We used values obtained by fitting *in vivo* voltage clamp experiments that account for the imperfect voltage-clamping caused by the powerful voltage-dependent conductances [24].

The fast delayed rectifier (FDR) has conductance 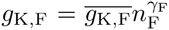 and the slow delayed rectifier (SDR) has conductance 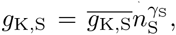 where *γ*F and *γ*S are the number of gating particles; 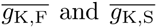 are their maximum conductances. The HH activation gating variables, *n*_F_ and *n*_S_, change with time following the differential equations:

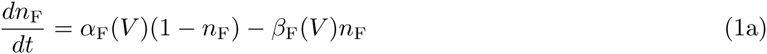

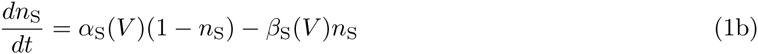

where the parameters *α*_F_, *α*_S_, *β*_F_ and *β*_S_ are the activation and deactivation rates respectively. The activation and deactivation rates depend on voltage, as given by the following formulae:

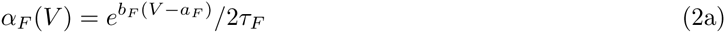

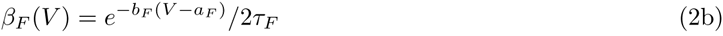

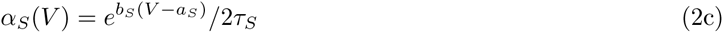

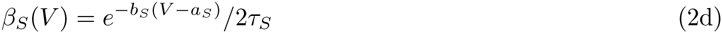

See Table 1 for the values of the parameters.

**Table 1:**
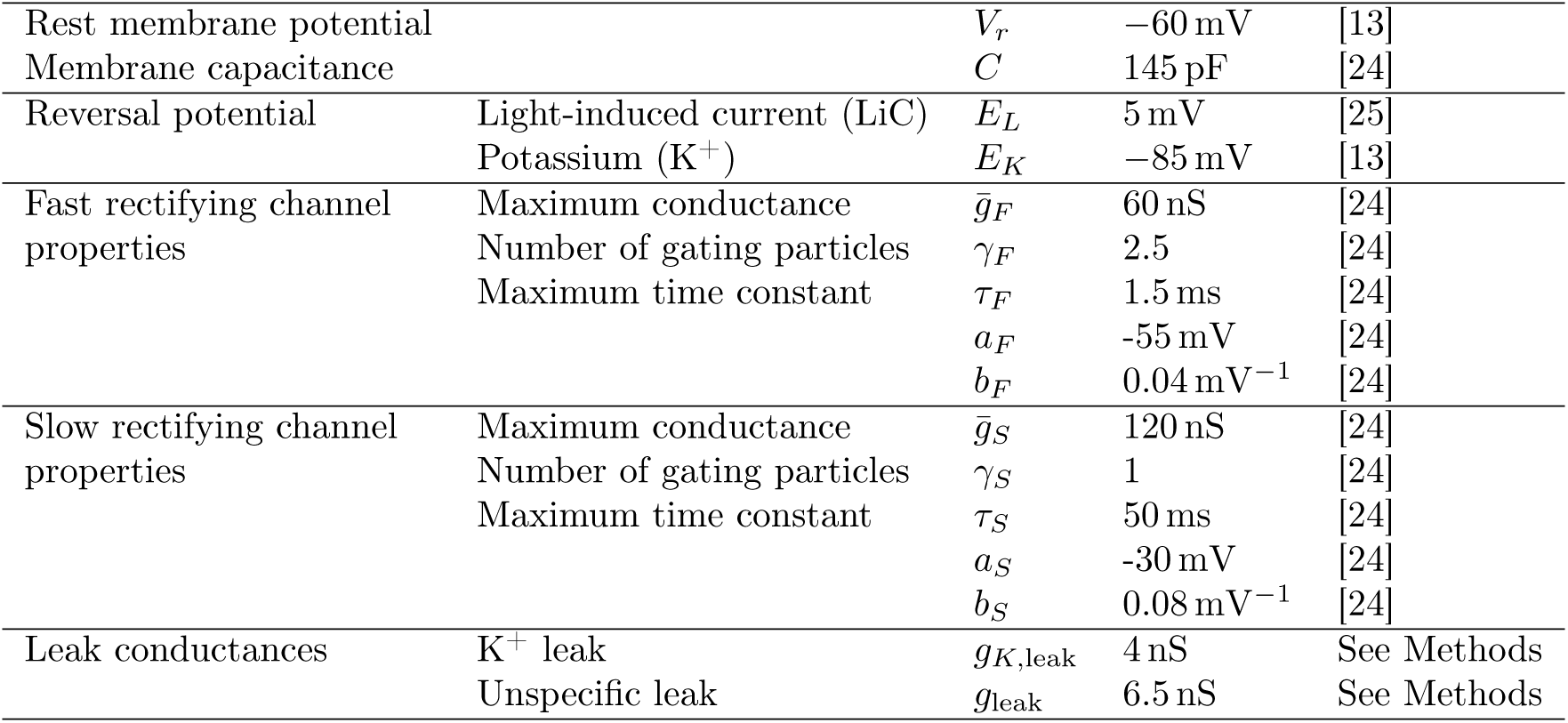
Biophysical parameters of the model blowfly R1-6 photoreceptor.

|  |  |  |  |  |
| --- | --- | --- | --- | --- |
| Rest membrane potential | | $V_r$ | −60 mV | [13] |
| Membrane capacitance | | $C$ | 145 pF | [24] |
| Reversal potential | Light-induced current (LiC) | $E_L$ | 5 mV | [25] |
| | Potassium ($K^+$) | $E_K$ | −85 mV | [13] |
| Fast rectifying channel properties | Maximum conductance | $\overline{g}_F$ | 60 nS | [24] |
| | Number of gating particles | $\gamma_F$ | 2.5 | [24] |
| | Maximum time constant | $\tau_F$ | 1.5 ms | [24] |
| | | $a_F$ | −55 mV | [24] |
| | | $b_F$ | 0.04 mV<sup>−1</sup> | [24] |
| Slow rectifying channel properties | Maximum conductance | $\overline{g}_S$ | 120 nS | [24] |
| | Number of gating particles | $\gamma_S$ | 1 | [24] |
| | Maximum time constant | $\tau_S$ | 50 ms | [24] |
| | | $a_S$ | −30 mV | [24] |
| | | $b_S$ | 0.08 mV<sup>−1</sup> | [24] |
| Leak conductances | $K^+$ leak | $g_{K,leak}$ | 4 nS | See Methods |
| | Unspecific leak | $g_{leak}$ | 6.5 nS | See Methods |

The DR properties are comparable with previous experimental results on isolated photoreceptors, after accounting for a 15 mV to 20 mV negative shift of the membrane voltage [13, 26]. The maximum total K^+^ conductance is 184 nS, with the activation curves of the SDR and FDR separated by 11 mV. These values are within the estimated ranges of 100 nS to 200 nS and 10 mV to 15 mV reported by [13]. The maximum time constants for FDR and SDR (*τ*_F_ = 1.5 ms and *τ*_S_ = 50 ms, respectively) are also consistent with previously published experimental values (1 ms to 10 ms and 5 ms to 40 ms, respectively) [13].

### 2.2 A biophysical model of the blowfly R1-6 photoreceptor membrane

Our model of the blowfly R1-6 photoreceptor membrane incorporates both types of K^+^ conductances (FDR and SDR, Figure 1a,b); and the Na^+^/K^+^ ATPases, molecular pumps with very slow dynamics [27, 28, 26], which we treat as a fixed current source equilibrating the steady-state K^+^ current during small perturbations around a particular depolarisation (Figure 1b).

**Figure 1:**
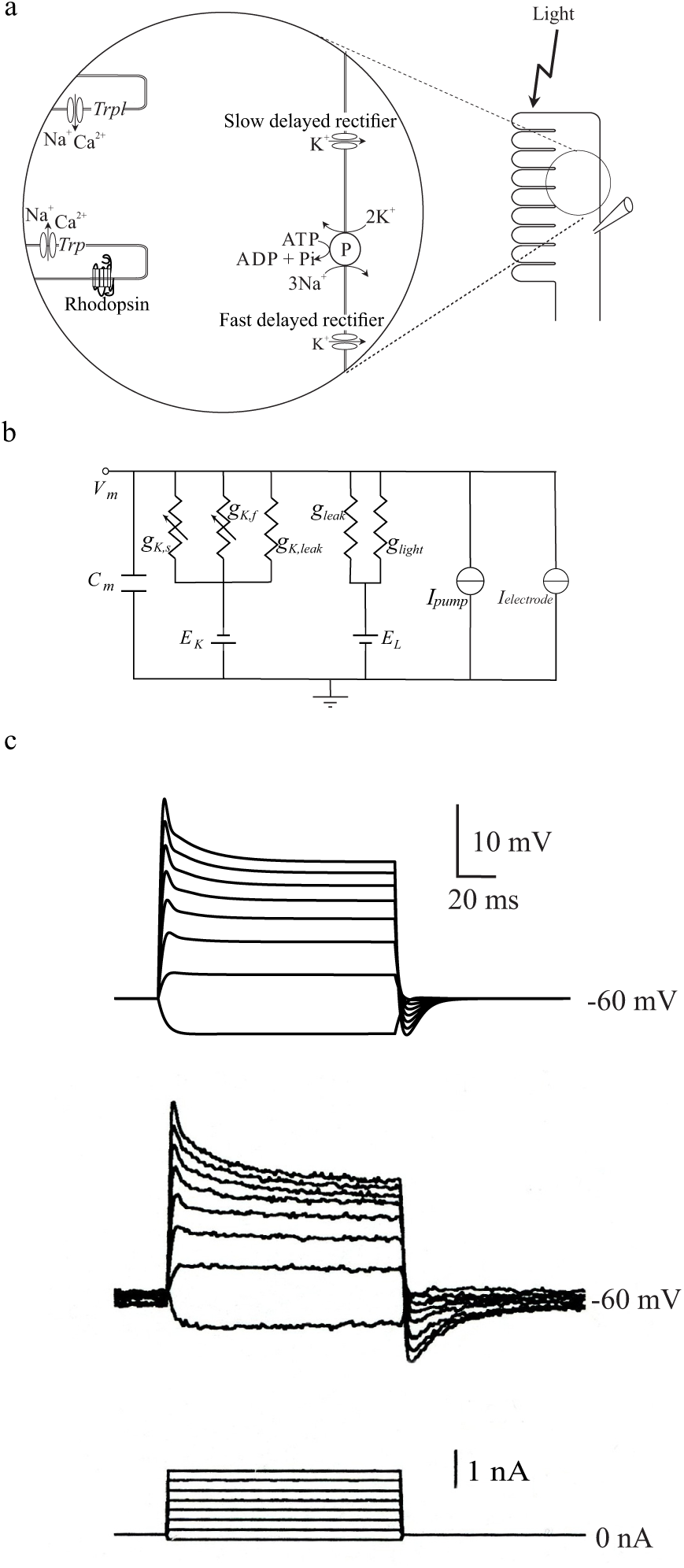
A Hodgkin-Huxley type model reproduces the voltage responses of a blowfly R1-6 photoreceptor. (a) A schematic diagram of a photoreceptor showing the major components that generate voltage responses to light. (b) A literal electrical circuit model of the photoreceptor membrane incorporating the capacitance (*C*), the light-induced conductance (*g*_light_), voltage-independent leak conductances (*g*_leak_ and *g*_K,leak_), the fast and slow delayed rectifier conductances (*g*_K,F_ and *g*_K,S_), and the reversal potentials (*E*_L_ and *E*_K_). The Na^+^/K^+^ ATPase generates the pump current (*I*_P_). (c) Comparison of the voltage responses of the literal electric circuit model with those of a blowfly photoreceptor recorded *in vivo*. (*Upper panel*). The voltage responses of the model blowfly R1-6 photoreceptor to injected current. (*Middle panel*). The intracellularly recorded voltage responses of a blowfly R1-6 photoreceptor to the same set of injected current pulses. R1-6 voltage response reproduced from [24]. (*Lower panel*). The current pulses injected to produce the voltage responses.

The model has a membrane capacitance of 145 pF, estimated during hyperpolarization to negative holding voltages, when all the K^+^ conductances are closed [24]. The K^+^ reversal potential of blowfly photoreceptors is −85 mV [13]. The reversal potential of the LiC in the model is 5 mV, the same as that of fruit fly R1-6 photoreceptors [25].

The model also includes a K^+^ leak conductance, *g*_K,leak_, of 4 nS to reproduce the behaviour of hyperpolarised photoreceptors *in vivo* [24]. An additional leak conductance *g*_leak_ of unknown nature and origin is needed to depolarise the photoreceptor to the resting membrane potential from the K^+^ reversal potential. We assume this leak conductance has the same reversal potential as the LiC [13, 29]; with a value of 6.5 nS, to reproduce the dark-adapted resting potential of −60 mV. The model does not incorporate the photoreceptor axon, the effect of which will be small because its input resistance (∼100 MΩ [30]) is much greater than the membrane impedance at all light intensities.

### 2.3 Obtaining the steady-state currents

The steady-state values of the conductances at different membrane potentials were calculated using well-known procedures (e.g. [29]). Each steady-state light intensity generates a light-induced conductance, *g*_light_, and the corresponding depolarisation a total K^+^ conductance, *g*_K_. The K^+^ and light-induced currents flowing through the conductances are given by:

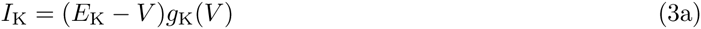

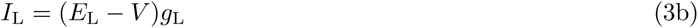

where *g*_K_(*V*) = *g*_K,F_(*V*) + *g*_K,S_(*V*) + *g*_K,leak_ is the total K^+^ conductance and *g*_L_ is the total depolarising conductance, the sum of both the leak conductance and the light-induced conductance *g*_L_ = *g*_light_ + *g*_leak_.

The Na^+^/K^+^ ATPase maintains the ionic concentration in the photoreceptor by pumping 2 K^+^ ions in and 3 Na^+^ ions out in each cycle [27]. Consequently, maintaining a constant internal K^+^ concentration produces a net pump current, *I*_P_, of the same sign and half the magnitude of the K^+^ current [28, 31]:

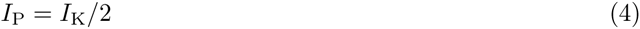

In the steady state, the currents across the membrane must sum to zero:

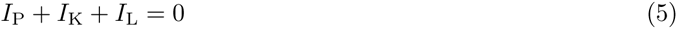

a condition that, together with Equations 3 and 4, gives the following equation:

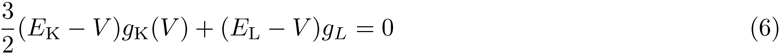

Because the K^+^ conductance is known, we can solve Equation 6 to obtain the total depolarizing conductance, *g*_L_. We first solve it for the dark-adapted photoreceptor to obtain *g*_leak_, and then for the light-adapted photoreceptor to obtain *g*_light_ = *g*_L_ − *g*_leak_.

### 2.4 Current injection simulations

We drove the photoreceptor membrane model with current from a given steady-state voltage, set by an increase in light-induced conductance, *g*_light_. The time step of integration was 0.025 ms. We numerically integrated the differential equations using a forward Euler integrator. The model reproduces the voltage response to injection of step current pulses (Figure 1c).

### 2.5 A closed formula for the membrane impedance

The response of a voltage-dependent conductance to small perturbations around a given voltage can be approximated by an RrL electrical circuit [32]. For a DR with conductance, *g* = *g̅n*^γ^, it is formed by a resistance, *R*, connected in parallel with a phenomenological branch containing a resistor, *r*, and an inductor, *L*, whose values are given by

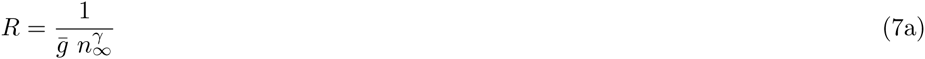

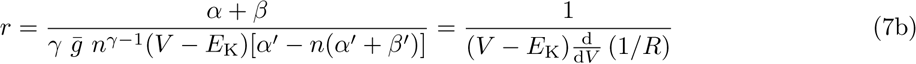

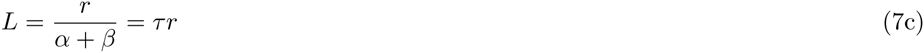

where *α* = *α*(*V*), *β* = *β*(*V*) are, as before, the activation rate, deactivation rate of the HH variable *m* at the voltage *V*; *τ* = 1/(*α* + *β*) and *n*_*∞*_ = *α*/(*α* + *β*) are respectively the time constant and the steady state value of *m* at the voltage *V*; and a prime (‘) represents the derivative with respect to voltage.

We then use *R, r* and *L* to compute the impedance of each DR:

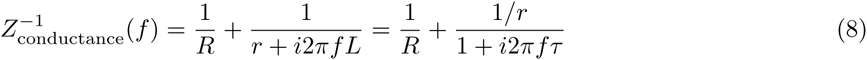

The electrical response of the photoreceptor membrane for small signals around a certain voltage can be approximated by the circuit formed by the light-induced and leak conductances, the RrL equivalent circuit of the two K^+^ conductances and the membrane capacitance [21] (Figure 2). The resulting impedance is

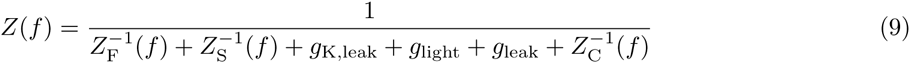

where *Z*_F_(*f*) and *Z*_S_(*f*) are the impedances of the fast and slow rectifiers respectively (calculated using Equation 8), and *Z*_C_(*f*) = 1/*i*2*πfC* is the impedance of the membrane capacitance.

**Figure 2:**
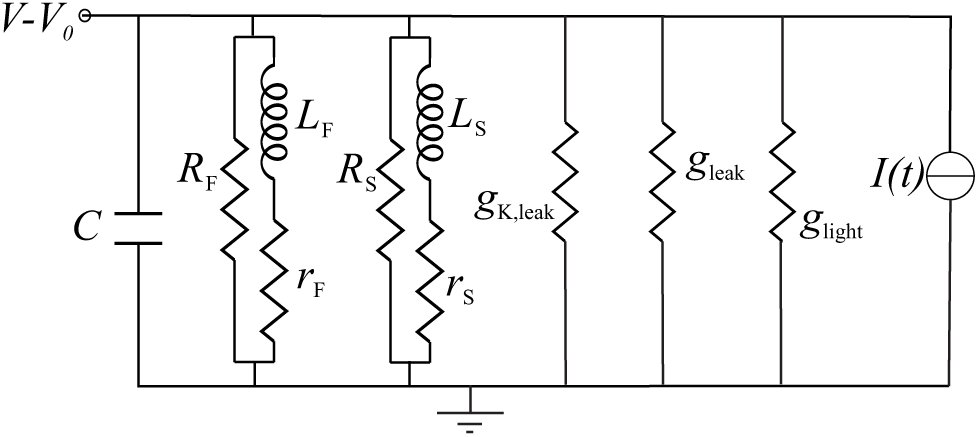
A phenomenological electrical circuit (RrLC circuit) of the photoreceptor describing the small signal behaviour of the membrane to injected currents around a depolarised voltage, *V*_0_. In this circuit, the delayed rectifiers *g*_K,S_ and *g*_K,F_ are each modelled as a resistance, *R*, in parallel with a branch composed by a resistance, *r*, and an inductance, *L*, in series.

### 2.6 Input resistance and membrane resistance

At each voltage, the input resistance of the photoreceptor membrane is defined as the low frequency limit of the impedance:

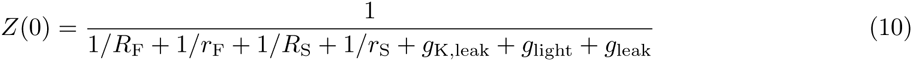

It differs from the membrane resistance, defined as the inverse of the sum of all conductances, both voltage-dependent and voltage-independent, which is:

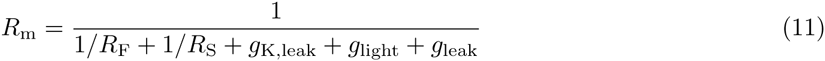

### 2.7 Energy consumption

We obtain the net pump current from Equation 4. Because the Na^+^/K^+^ ATPase hydrolyses an ATP molecule for every 2 K^+^ ions pumped in and 3 Na^+^ ions pumped out [28], we can calculate the pump energy consumption:

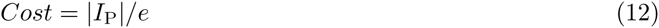

in hydrolysed ATP molecules/s when the current, *I*_P_, is given in amperes and the elementary charge, *e*, is in coulombs.

## 3 Results

We used a Hodgkin-Huxley type (HH) membrane model [23] to determine the effects of voltage-dependent K^+^ conductances upon the gain (i.e. maximum impedance), bandwidth and energy consumption of the blowfly R1-6 photoreceptor membrane. The model incorporated an inward light-induced conductance, opposed by two outward voltage-dependent K^+^ conductances; a fast delayed rectifier (FDR) and a slow delayed rectifier (SDR) (see Methods; Figure 1a,b) [13, 33, 24].

### 3.1 FDR and SDR shape the voltage response to injected current

The voltage responses of the model to injected current compare well with those recorded *in vivo* (Figure 1c) suggesting that the model’s parameters are appropriate. When the model is at the dark resting potential, the light-induced conductance is zero and the membrane’s responses to pulses of positive and negative current exhibit the rectification that is the hallmark of DRs (Figure 1c, 3a). Positive current pulses activate the DRs, thereby reducing the extent of depolarization. Negative current pulses have the opposite effect. Consequently, both depolarising and hyperpolarising voltage responses deviate from the RC charging curve of a passive model membrane with the same resistance at rest (Figure 1c, 3a-c). In most cases the response of the passive model to a given current reaches its maximum amplitude considerably later than the active model’s response, the exception being large hyperpolarisations that drive the membrane well beyond its physiological voltage range.

**Figure 3:**
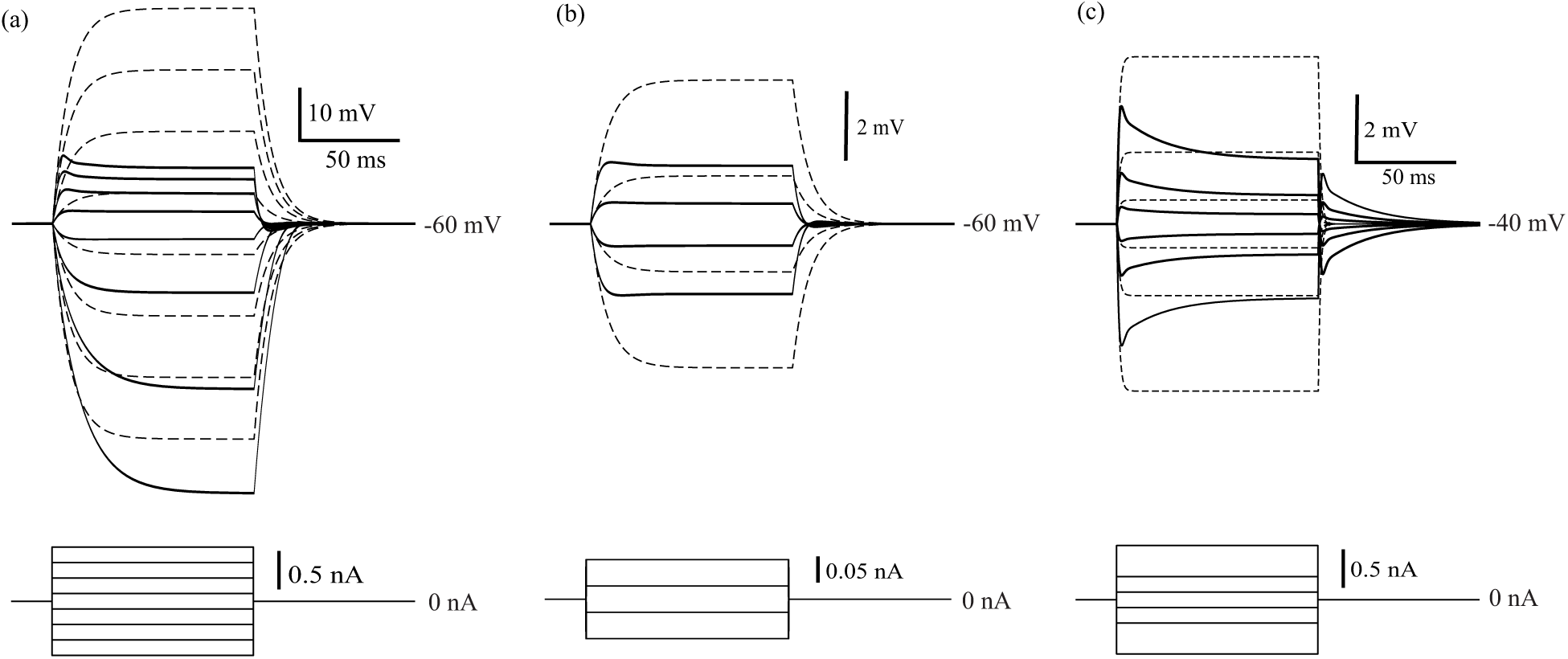
Voltage responses of the Hodgkin-Huxley type model to large and small current pulses. (a) (*Upper panel*). Voltage responses evoked by large injected current pulses into the photoreceptor membrane model with active DRs (black, solid) or into the passive photoreceptor membrane model (black, dashed) in dark-adapted conditions. (*Lower panel*). The set of current pulses that evoked the voltage responses. (b) (*Upper panel*). Voltage responses evoked by small injected current pulses into the photoreceptor with active DRs (black, solid) or the passive photoreceptor (black, dashed) at rest. (*Lower panel*). The set of current pulses that evoked the voltage responses. (c) (*Upper panel*). At high light intensities that depolarise the photoreceptor to −40 mV, injected current pulses evoke small voltage responses from the active photoreceptor containing voltage-dependent K^+^ conductances (black, solid) and from a passive photoreceptor (black, dashed). (*Lower panel*). The set of current pulses that evoked the voltage responses.

The waveform of the model membrane’s response to injected current changes with amplitude because the SDR’s voltage activation range is 11 mV more positive than that of the FDR [13, 24]. Small depolarisations almost exclusively activate the FDR and, because its activation time constant is short, the membrane potential rapidly equilibrates, resulting in a step-like voltage response. Larger depolarisations activate the SDR, producing a pronounced sag in response that reflects the SDR’s longer activation time constant.

The small responses evoked by small currents (≤ 0.1 nA) are symmetrical: The response to a small negative step current is equal and opposite to the response to a positive step current of identical amplitude (Figure 3c). This linear behaviour does not, however, imply that the effects of voltage-dependent conductances are negligible. A direct comparison of the responses of the active and passive membranes shows that the DRs are still reducing maximum response amplitude and time-to-peak. Indeed, these small responses (≤ 2 mV) lie within the range where the membrane behaves linearly, according to a well-established model that uses active components, namely inductances [23, 21].

Depolarising the model membrane to a steady level with light-induced current (LiC), as happens when a photoreceptor is exposed to a constant background light, has three effects on the response to current pulses: (i) it reduces the response amplitude; (ii) it reduces rectification; and (iii) it increases response sag. The response amplitude is reduced because the membrane has a lower resistance, produced by the increase in both the light-induced conductance and the K^+^ conductance. Rectification is reduced because placing the Ohmic light-induced conductance in parallel with the DRs reduces the voltage-dependence of the membrane’s total conductance. This buffering by a parallel fixed conductance increases the voltage range over which the membrane behaves linearly. The sag is prominent in all responses, even the smallest, because the SDR is activated at higher levels of depolarization.

As observed at the dark resting potential, the voltage responses to injected current initially follow the trajectory of the RC charging curve of an equivalent passive membrane (dashed line) but deviate after several milliseconds (Figure 3) due to the activation or deactivation of the two DRs. Because the membrane potential is the result of a balance between opposing currents, a change in DR conductance produces negative feedback, with a delay due to the finite time needed for activation or deactivation [13, 33, 24]. Thus, there are two effects of the negative feedback on the linear voltage responses to injected current. First, negative feedback decreases the voltage response to injected current, reducing the amplitude below that expected from Ohms law (Figure 3b,c). Second, negative feedback truncates the step response, reducing the time-to-peak (Figure 3b,c), thereby increasing the frequency response of the membrane.

### 3.2 FDR and SDR shape the model membrane’s frequency response

To determine the effect of DRs on the model membrane’s frequency response, we calculated the membrane’s impedance, *Z*(*f*), which is the amplitude of the voltage response, *V* (*f*), divided by the amplitude of the current producing it, *I*(*f*), at the corresponding frequency.

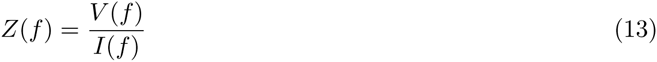

Note that *Z*(*f*), for each frequency, *f*, is a complex number whose modulus corresponds to the gain with which the membrane converts a modulation in current, especially LiC, into a modulation of voltage.

We calculated *Z*(*f*) using a closed formula, solving an RrLC electrical circuit that represents the linearisation of the membrane with voltage-dependent conductances (see Methods, Figure 2). The impedance calculated with the RrLC circuit formula is identical to that determined from the response of the HH model to low variance white noise current [21].

In the absence of voltage-dependent conductances, the membrane is a passive, RC low-pass filter (Figure 4a) whose impedance, *Z*_RC_(*f*) is determined by its resistance, *R*_m_, and its capacitance *C*:

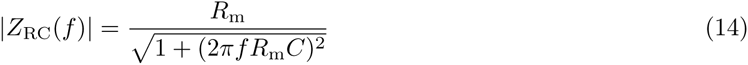

**Figure 4:**
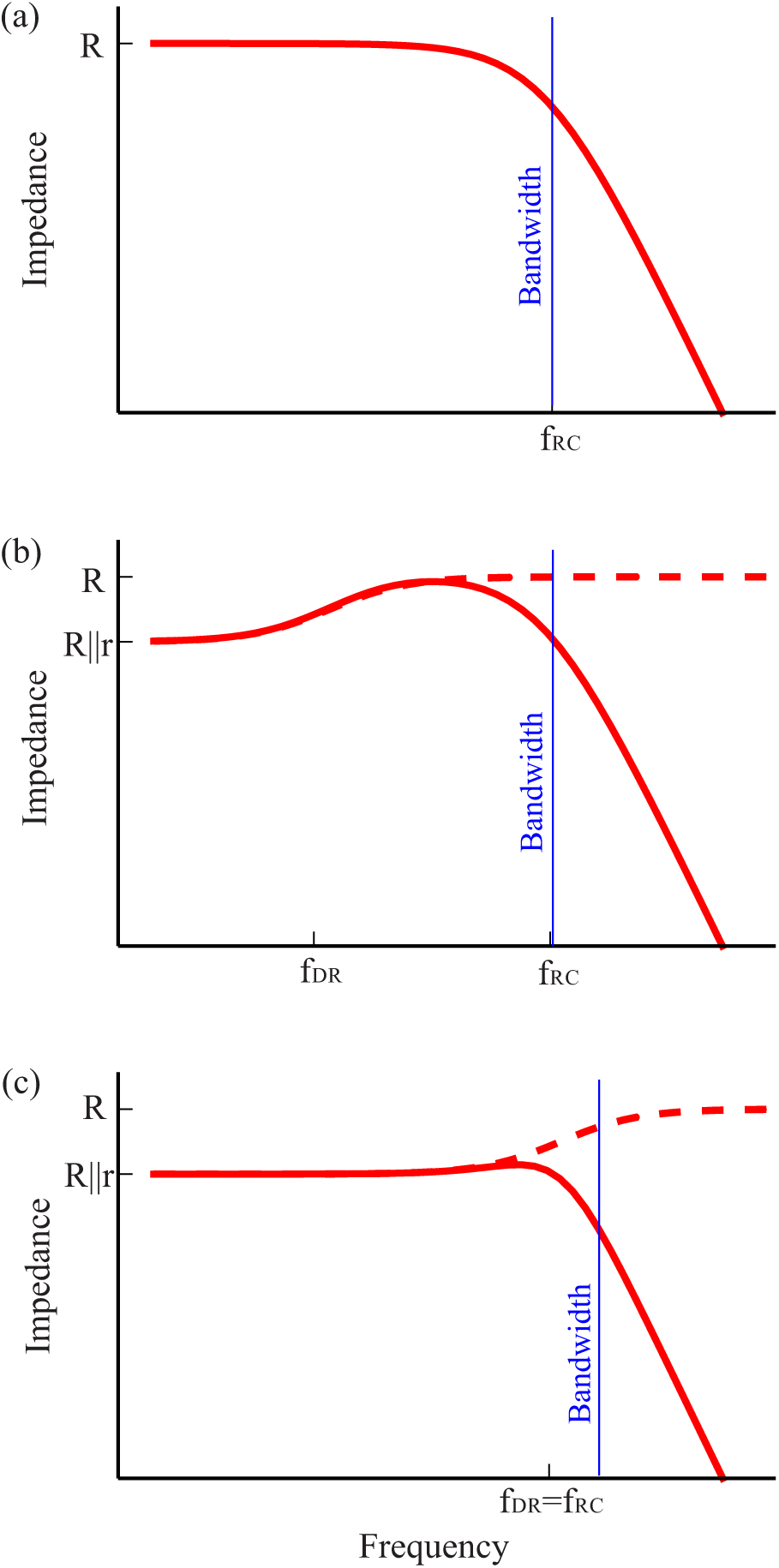
Voltage-dependent K^+^ conductances tune the membrane impedance. (a) The impedance of a passive membrane without voltage-dependent K^+^ conductances, in a log-log plot. The cut-off frequency of the passive membrane, *f*_RC_, which coincides with the bandwidth, and the membrane resistance, *R*_m_ = *R*, are indicated. (b) Impedance of an active photoreceptor membrane with a single voltage-dependent K^+^ conductance including (solid) or excluding (dashed) the membrane capacitance, in a log-log plot. The transition frequency of the conductance, *f*_DR_, is lower than the cut-off frequency of the passive membrane, *f*_RC_. Impedance at low frequencies decreases to *R‖r* due to negative feedback, which in the RrLC model is captured by the phenomenological branch with a resistor, *r*, and the resulting membrane bandwidth is only slightly higher than *f*_RC_. (c) As in b but with a single voltage-dependent K^+^ conductance, the transition frequency of which coincides with the cut-off frequency of the membrane. The impedance of the membrane decreases towards *R‖r* and the resulting membrane bandwidth is higher than *f*_RC_.

The bandwidth (3dB cut-off frequency) of this passive membrane is *f*_RC_ = 1/2_*π*_*R*_m_*C*. Note that in a passive membrane, the input resistance (the low frequency limit of the impedance) has the same value as the membrane resistance, *Z*(0) = *R*_m_.

Incorporating a DR produces negative feedback that attenuates frequencies below its transition frequency, *f*_DR_ = 1/2_*πτ*_, where *τ* is the activation time constant of the DR. This can be understood by considering the equivalent RrL circuit of the DR, consisting of a resistance, *R*, connected in parallel to a phenomenological branch composed by a resistance, *r*, and an inductance, *L*. Below the transition frequency, the inductance acts as a short circuit, and the impedance of the DR corresponds to *R* and *r* in parallel, *R‖r* = (*R*^−1^+*r*^−1^)^−1^. Above the transition frequency, the impedance of *L* increases, effectively opening the circuit at the rL branch and producing an impedance of *R* (Figure 4b,c). Unlike in the passive membrane, in this active membrane the input resistance is smaller than the membrane resistance, *Z*(0) < *R*_m_(Figure 4).

Together with the capacitive roll-off of the membrane, a DR with a transition frequency well below the passive membrane cut-off frequency, *f*_DR_ τ *f*_RC_, produces a band-pass filter peaking at the membrane resistance, *R*_m_ = *R* (Figure 4b). However, if the transition frequency is close to or higher than the passive membrane cut-off frequency, the impedance is reduced to *R‖r* below the capacitive roll-off (Figure 4c). In the latter case, the membrane acts as a low-pass filter but its input resistance, *Z*(0) = *R‖r*, is still lower than the membrane resistance, *R*_m_ = *R*.

Having treated a simple case, the effects of single DR, we now consider the blowfly photoreceptor membrane model, by incorporating the FDR and the SDR, together with the leak and light-induced conductances (Figure 1b, 2). This membrane’s impedance can be similarly analysed in terms of the contribution of both the FDR and SDR (Equation 9, Figure 5a). The DRs differ in their activation time constants across the physiological range of membrane potentials; changing upon light adaptation from 1.5 ms to 1 ms for the FDR and from 10 ms to 40 ms for the SDR [24]. Consequently, the SDR reduces the impedance only at low frequencies, producing band-passing (cf. Figure 4b), whereas the FDR reduces the impedance across all frequencies (cf. Figure 4c).

**Figure 5:**
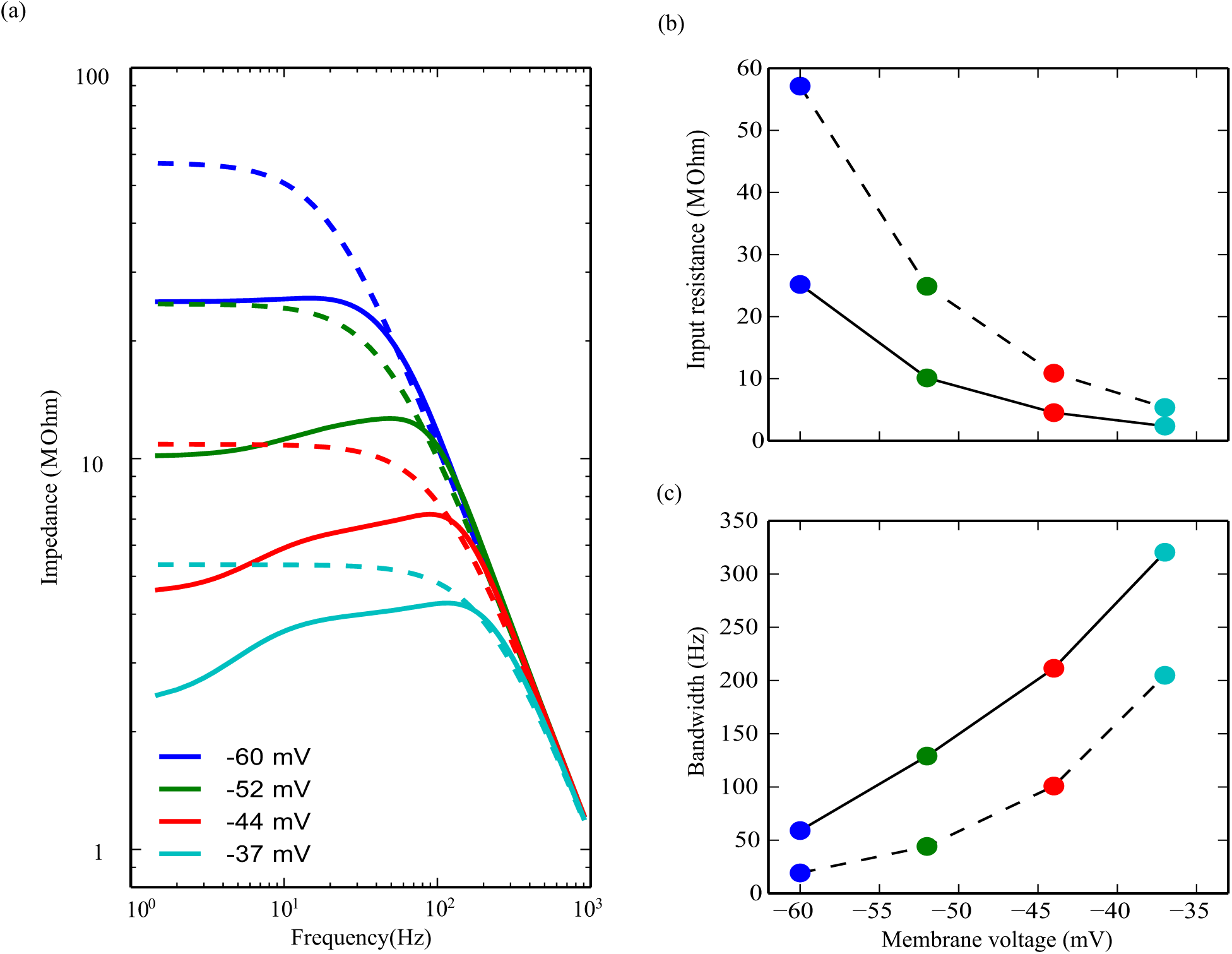
Negative feedback from voltage-dependent K^+^ conductances reduces photoreceptor membrane impedance and increases bandwidth. (a) Photoreceptor impedance in the dark and when depolarised by light to three different membrane potentials (solid) compared to passive RC membranes with the same resistance and capacitance (dashed). (b) The low frequency limit of the active photoreceptor impedance, i.e. the input resistance, (solid) compared to that of the passive membrane with the same resistance (dashed) at different membrane potentials. Note than in the passive membrane input resistance equals membrane resistance, so the dashed line also represents the membrane resistance of both the active and the passive photoreceptor. (c) The active photoreceptor membrane bandwidth, i.e. 3dB cut-off frequency of the impedance, compared to the bandwidth of the passive membranes with the same resistance (dashed) at four different membrane potentials.

Due to the FDR, then, the impedance is lower than the membrane resistance across the photoreceptors’ physiological membrane voltage range (Figure 5a). This is a consequence of the activation time constant of the FDR, *τ*_F_(*V*), being fast enough to lower all frequencies below the capacitive roll-off. At the dark resting potential of −60 mV and at dim light intensities, the membrane impedance acts essentially as a low-pass filter for the LiC but with an input resistance that is lower than the membrane resistance (Figure 5a). At higher membrane potentials (from −44 mV to −36 mV), the band-passing becomes more pronounced because the SDR conductance is activated more strongly, depressing the impedance below 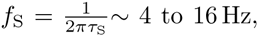 where *τ*_S_(*V*) is activation time constant of the SDR (Figure 5a). Thus, the RrLC model illustrates how the membrane’s impedance is tuned by voltage-dependent conductances according to the level of depolarization, as is observed in experiments in blowfly photoreceptors [13, 34, 24].

We compared the properties of the active (i.e. voltage-dependent) membrane to a passive RC membrane with identical resistance and capacitance, to quantify the effects of the negative feedback produced by the DRs (Figure 5a-c). We calculated the input resistance and bandwidth of both membranes at all light intensities. Due to negative feedback from the FDR, the active membrane has an input resistance of 24.7 MΩ in the dark. The input resistance of the passive RC membrane, which is identical to its membrane resistance of 56.1 MΩ, significantly exceeds the input resistance of the active membrane (Figure 5b). The active membrane bandwidth in the dark is 59 Hz (Figure 5c), considerably higher than that of a passive membrane with the same resistance and capacitance, 19 Hz.

At low light intensities, when the membrane voltage is −52 mV, the negative feedback lowers the input resistance of the active membrane to 10.1 MΩ, well below the membrane resistance of 24.9 MΩ (Figure 5b). The bandwidth of the active membrane increases to 129 Hz, larger than that of a passive membrane (44 Hz). At the highest light intensities, when the LiC depolarises the membrane to −37 mV and both the SDR and FDR are strongly activated, the input resistance of the active membrane drops to 2.4 MΩ, again lower than the membrane resistance of 5.4 MΩ (Figure 5b). The active membrane bandwidth of 320 Hz exceeds that of a passive membrane with the same resistance and capacitance (205 Hz) by an even greater margin.

Thus, by decreasing input resistance 10-fold upon depolarisation, the photoreceptor operates in a high-impedance, low-bandwidth regime at low light intensities shifting to a low-impedance, high-bandwidth regime at high light intensities [13, 24] (Figure 5). At all light intensities, the negative feedback produced by the DRs further increases bandwidth and decreases the membrane impedance, particularly at low frequencies.

### 3.3 DRs reduce the energy cost of membrane bandwidth

The increase in photoreceptor membrane bandwidth produced by light-induced depolarisation is associated with an increase in energy consumption. The light-induced conductance increases to depolarize the photoreceptor, thereby increasing the K^+^ current by increasing the driving force on K^+^ ions and by activating DRs. The flux of ions through these conductances is reversed by the Na^+^/K^+^ pump, which uses energy from ATP hydrolysis to maintain ionic concentration gradients [27, 28]. The rate of ATP consumption is specified by the pump current, which in the steady state is half the K^+^ current (Equation 4). Our HH type membrane model predicts that when the photoreceptor membrane is depolarised to the steady state level recorded in bright light (-37 mV), the pump current is 1.65 nA, more than ten times its value in the dark. This translates into an increase in energy consumption from 8.7 × 10^8^ ATP/s to 1.03 × 10^10^ ATP/s.

We used our HH type model of the photoreceptor membrane (Figure 1b) to calculate the effects of voltage-dependent K^+^ conductances on energy consumption by the Na^+^/K^+^ pump. We compared the model with FDR and SDR operating at a given membrane potential to a passive RC membrane with the same capacitance and bandwidth operating at the same membrane potential (Figure 6a-c). This matched passive membrane model was constructed by replacing the DRs in the HH type model with a fixed K^+^ conductance and adjusting its value, together that of the light-induced conductance, to achieve the same membrane potential and bandwidth.

**Figure 6:**
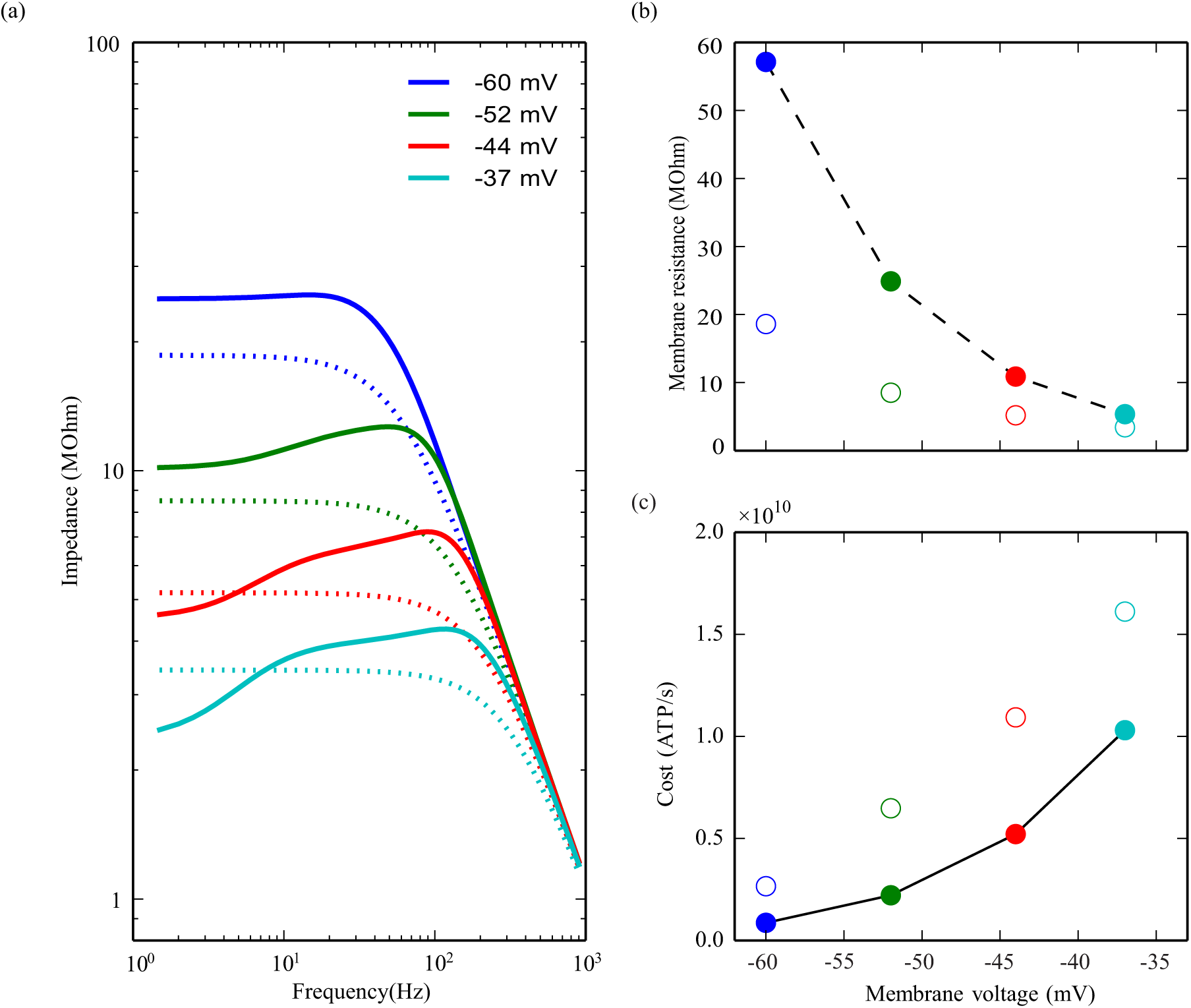
Voltage-dependent K^+^ conductances reduce the energy cost of achieving a particular bandwidth. (a) Photoreceptor impedance when depolarised by light to four different membrane potentials (solid) compared to that of matched passive membranes with the same capacitance and bandwidth (dashed). (b) The membrane resistance of the active membrane (dashed, filled circles) compared to that of the matched passive membranes (empty circles) at the four different membrane potentials considered. (c) The energy consumption (ATP molecules per second) of the active photoreceptor (solid, filled circles) compared to that of photoreceptors with matched passive membranes at different membrane potentials (empty circles).

At rest in the dark, and over the physiological range of steady-state depolarizations generated by background light, a matched passive membrane has a higher K^+^ conductance and light-induced conductance than its active counterpart, and hence a lower impedance and membrane resistance (Figure 6b), resulting in a higher energy consumption (Figure 6c). The differences are large. The passive membrane matched to the active membrane at the dark resting potential (-60 mV) has a resistance of 18.7 MΩ while that of the active membranes is 56.1 MΩ, three-fold higher. The passive membrane matched to the active membrane depolarised by bright light to −37 mV has a resistance of 3.4 MΩ, while the active membrane has a resistance of 5.4 MΩ, 58% higher (Figure 6c). At the dark resting potential, the energy consumption of the matched passive membrane is three-fold higher than that of the active membrane (2.6 × 10^9^ ATP/s versus 0.87 × 10^9^ ATP/s), and at the highest light levels 56% higher (1.6 × 10^10^ ATP/s versus 1.0 × 10^10^ ATP/s) (Figure 6c). These rates of ATP consumption demonstrate that the negative feedback provided by voltage-dependent K^+^ conductances makes large energy savings across the physiological range of mean membrane potentials by reducing the conductance required to achieve a given membrane bandwidth.

By activating progressively across the membrane’s voltage operating range a voltage-dependent K^+^ conductance can also save energy [22], but such savings have not been quantified. To estimate these savings, we assessed the performance of each matched passive membrane across the range of depolarizations produced by background light (Figure 7a-c). This was achieved by changing the light-induced conductance of each matched passive membrane. As expected [13], the range of bandwidths achieved by a passive membrane was smaller than the range achieved by the active membrane (Figure 7a). For example, the passive membrane matched at the dark resting potential (the “dark” passive membrane) has a bandwidth of 100 Hz at −37 mV, 70% less than that of the active membrane, and too low to transmit the high frequencies seen by blowflies in bright light. Conversely, the passive membrane matched to the active membrane in bright light at −37 mV (the “bright-light” passive membrane) has a more than three-fold greater bandwidth than the active membrane at the dark resting potential. This greater bandwidth is not useful at low light levels because high frequency signals are lost in photon noise [35, 36]. Excessive bandwidth comes at a high cost. At the dark resting potential the “bright-light” passive membrane consumes ten times more energy than the active membrane (Figure 7c). This is almost as much energy as the active membrane consumes in bright light at −37 mV. Thus, the ability of the active membrane to reduce its K^+^ conductance with membrane potential substantially reduces energy costs.

**Figure 7:**
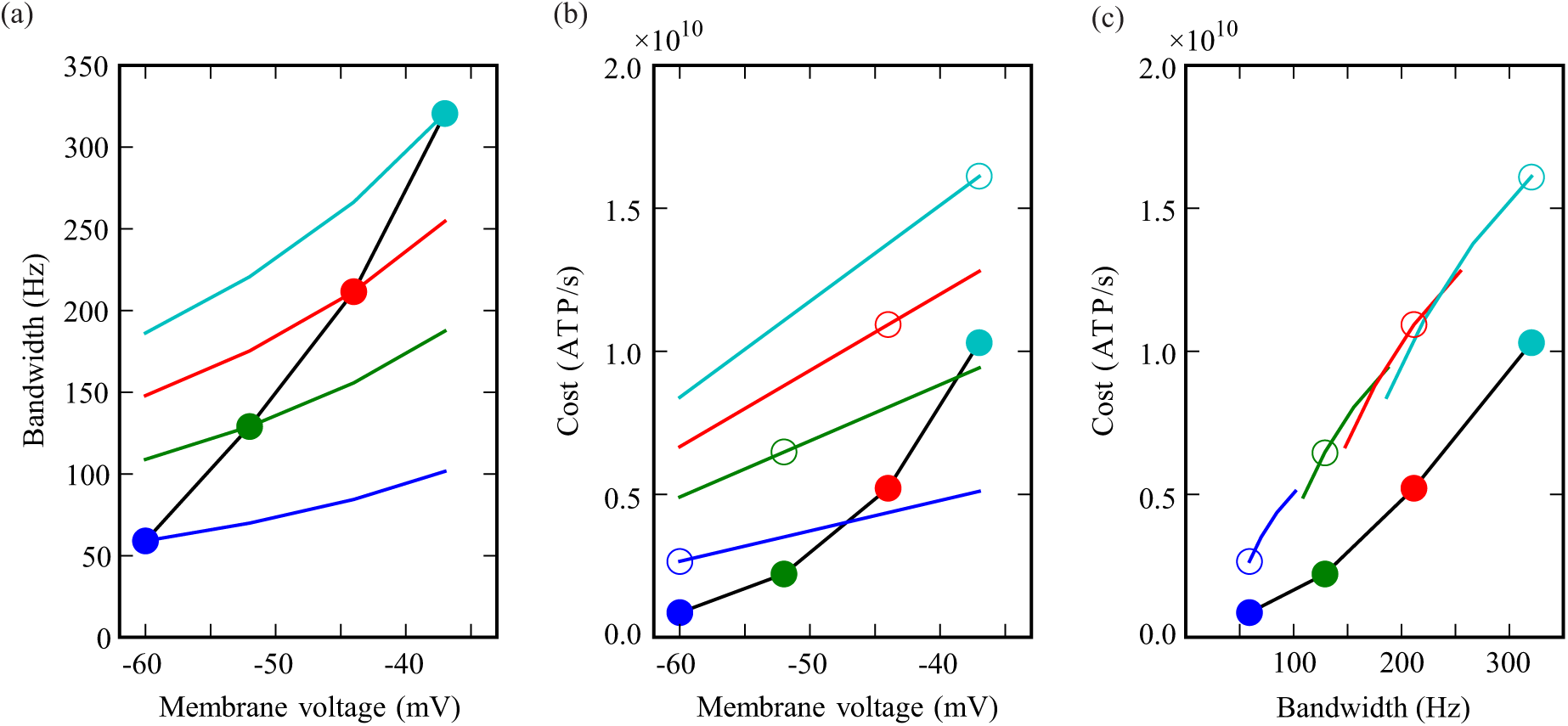
Voltage-dependent K^+^ conductances improve the energy efficiency of signalling and permit differential energy investment. (a) Bandwidth (3dB cut-off frequency of the impedance) of active photoreceptor membrane (black line with filled circles) compared to the bandwidth of the photoreceptors with passive membranes considered in Figure 6, which are then depolarised by increasing the light-induced conductance (coloured lines). Each line corresponds to a photoreceptor with passive membrane intersect the line corresponding to the photoreceptor with active membrane at the depolarisation where each passive membrane was chosen to have the same capacitance and bandwidth as the photoreceptor with active membrane. Then they diverge, showing that bandwidth increases less with depolarisation in the photoreceptors with passive membranes. (b) Energy consumption (ATP molecules per second) of the photoreceptor with active membrane (black line with filled circles) compared to that of passive matched membranes. Even at the depolarisation where the photoreceptors with passive membranes were matched in bandwidth, cost is larger in the passive photoreceptor (empty circles) than in the active photoreceptor (filled circles). The photoreceptors with passive membranes are driven to different steady state depolarizations by changing the light-induced conductance (coloured lines). (c) The energy (ATP molecules per second) that active and passive photoreceptors spend to achieve a given bandwidth. Photoreceptors with passive membranes (empty circles and coloured lines) consume more energy to achieve a particular bandwidth than the photoreceptor with active membrane (black line with filled circles).

## 4 Discussion

We modelled the membrane of blowfly R1-6 photoreceptors to determine how the activation of two voltage-dependent K^+^ conductances, the fast and slow delayed rectifiers (FDR and SDR), affects membrane impedance, bandwidth and energy consumption. We confirmed that when the membrane is depolarized by an increase in light-induced conductance, activation of the delayed rectifiers (DRs) decreases membrane impedance, and hence gain, and increases membrane bandwidth [13]. We also quantified the flow of K^+^ current and light-induced current (LiC) across the membrane and the energy consumed by the Na^+^/K^+^ pump, restoring the ions that crossed the membrane. These calculations enabled us to discover how voltage-dependent K^+^ conductances increase energy efficiency.

### 4.1 Delayed rectifiers increase efficiency by reducing the cost of membrane bandwidth

Increasing membrane bandwidth costs energy. A passive membrane is a low-pass RC filter whose bandwidth is inversely proportional to resistance. Consequently, ionic fluxes and the energy used by pumps to maintain ionic gradients increase in proportion to bandwidth. We find that in an active membrane voltage-dependent K^+^ conductances reduce the energy expended on bandwidth in two ways; by reducing the energy needed to achieve a particular bandwidth, and by adjusting bandwidth according to need (Figure 7).

The ability of voltage-dependent K^+^ conductances to reduce the energy cost of achieving a particular membrane bandwidth has not, to our knowledge, been reported before. Our modelling of the blowfly photoreceptor membrane demonstrates that this cost is reduced by two effects. Negative feedback lowers the amplitude of the voltage response to current; consequently the membrane’s impedance and input resistance are lower than its resistance. Impedance and input resistance determine how the membrane transforms current to voltage whereas membrane resistance determines ionic currents across the membrane, and hence energy consumption. For the photoreceptor membrane this means that two benefits of reducing impedance and input resistance, namely a lower gain and a higher bandwidth, are obtained with a higher membrane resistance at lower cost. The second effect, shunt peaking, is produced by the FDR acting as an inductance that boosts responses to higher frequencies, thereby increasing the gain bandwidth product (GBWP) [21]. Shunt peaking and the decreased impedance due to negative feedback are separable effects: when a DR’s activation time constant is substantially shorter than the membrane’s passive time constant, shunt peaking does not occur [21] but negative feedback still increases bandwidth by lowering input resistance.

Turning to the savings made by adjusting bandwidth according to need [22], our modelling demonstrates how they are made and estimates their magnitude. As the photoreceptor membrane is depolarised progressively by increasing light intensity, it shifts from a low-bandwidth, high-impedance (i.e. high-gain) membrane that consumes less energy to a high-bandwidth, low-impedance membrane that consumes more energy because of its lower resistance. This progressive increase in bandwidth contributes to the optimal filtering of natural images because, as light intensity increases, higher frequencies emerge from photon noise and are worth transmitting [35, 36]. By comparing the active photoreceptor membrane model containing voltage-dependent K^+^ conductances with passive models that contain voltage-independent K^+^ conductances, we find that voltage-dependent K^+^ conductances confer two advantages: they provide a greater increase in bandwidth upon light adaptation and they permit differential energy investment by adjusting bandwidth to signal quality (Figure 7). By avoiding investment in excessive bandwidth, energy consumption at low light levels is cut by 90% (Figure 7c). When one factors in the reductions in the cost of bandwidth brought about by negative feedback and shunt peaking the savings are substantial. Over the course of a summer’s day and night (14 hours light; 10 hours dark), our results suggest that DRs reduce the energy consumed by a blowfly photoreceptor membrane by almost 50%. Given that the high rate of daylight consumption accounts for more than 5% of the fly’s resting metabolic rate [37], this saving should increase a blowfly’s fitness.

### 4.2 Previous studies overestimated blowfly photoreceptor energy consumption and underestimated energy efficiency of information coding

Blowfly photoreceptor energy consumption has been estimated by applying experimental measurements of input resistance and membrane potential to a passive membrane model [31, 29]. The input resistance of the membrane which was measured by injecting current at a particular light intensity, was assumed to yield an approximation of the membrane resistance. We have shown, however, that voltage-dependent K^+^ conductances cause the input resistance to underestimate the membrane resistance by more than 50% at all light levels (Figure 5b). Consequently, previous studies [31, 29] overestimated photoreceptor energy consumption (Figure 8). In addition, our model shows that even taking the peak impedance as the membrane resistance, and incorporating it into a passive model, still overestimates the photoreceptor energy consumption (Figure 8, red dashed line).

**Figure 8:**
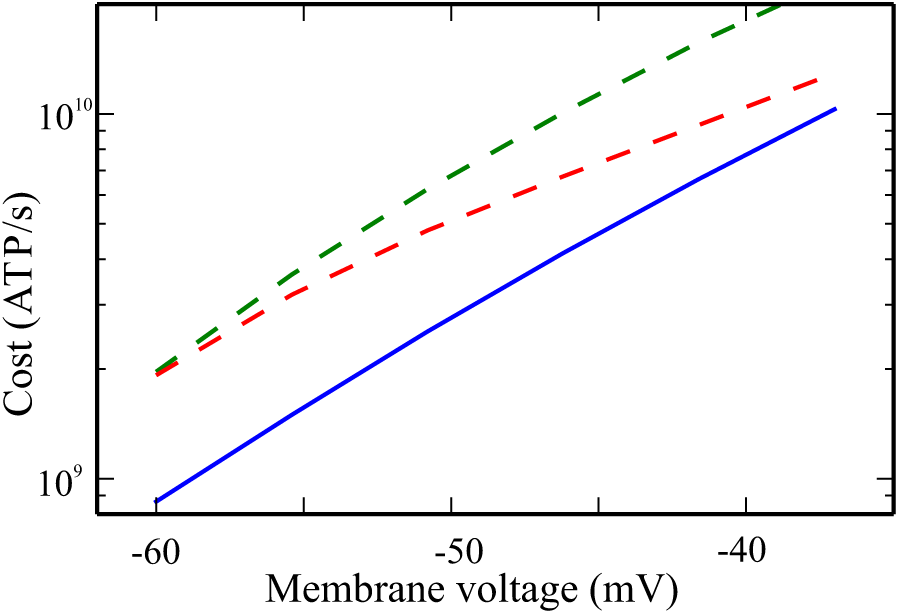
The energy consumption estimation of a blowfly R1-6 photoreceptor using impedance measurements depends on assumptions about membrane conductances. Energy consumption (ATP molecules per second) (blue, continuous) compared to the energy consumption that would be estimated either assuming that the input resistance (green, dashed) or the peak of the impedance (red, dashed) is the membrane resistance, and then modelling the photoreceptor as passive. Cost is plotted on a log scale to highlight relative estimation errors.

There is a second effect that could affect experimental estimations of blowfly photoreceptor energy consumption. Due to its internal resistance, the photoreceptor cell body will not be equipotential when driven by current injected at a single point along its length. Consequently, the impedance recorded by a sharp microelectrode overestimates the membrane impedance that filters the LiC, which is injected more uniformly through the microvilli arranged along the length of the photoreceptor cell body. By using a multi-compartment model, we show that this effect can be important at high light levels, when we predict an overestimate of the impedance of up to 50% (see Supplement 2). This effect will decrease, but not eliminate, the overestimation of energy consumption by previous studies (see Supplement 2).

The overestimation of energy consumption in previous studies led them to underestimate the energy efficiency of information coding. By combining the revised estimates of photoreceptor energy consumption from our model with experimentally-determined information rates [29] yields an energy efficiency of 1.7 × 10^−8^ bit/ATP at intermediate light intensities (10^3^ photons/photoreceptor/s), and of 6.5 × 10^−8^ bits/ATP at hight light intensities (10^6^ photons/photoreceptor/s). However, assuming that the input resistance is equivalent to the membrane resistance would underestimate energy efficiency as 6.9×10^−9^(or 7.8 × 10^−9^ bits/ATP, taking into account the effects of internal resistance). At high light intensities, a similar assumption would produce an estimate of photoreceptor energy efficiency (2.8 × 10^−8^ bits/ATP, or 4.2 × 10^−8^ bits/ATP if internal resistance is taken into account). Consequently, information coding is more energy-efficient at low and intermediate light levels than previously supposed.

Our estimates of photoreceptor energy consumption, made with a membrane model that includes the two DRs, compare well with experimental measurements of oxygen consumption from isolated blowfly retina [38]. Oxygen consumption measurements estimate photoreceptor energy consumption in the dark as 2.7 × 10^9^ ATP/s increasing to 6.5 × 10^9^ ATP/s when fully light-adapted [38]. Our equivalent estimates are 8.8 × 10^8^ ATP/s in the dark, increasing to 1.0 × 10^10^ ATP/s when fully light-adapted. Introducing the 25% inactivation of the DRs observed at high light levels [13] reduces the energy consumption of the fully light-adapted photoreceptor membrane model to 7.9 × 10^9^ ATP/s, bringing it closer to that estimated from retinal oxygen consumption [38].

The energy consumption of the second order interneurons in the blowfly visual system, the large monopolar cells (LMCs), has also been estimated from current injection [31]. Like photoreceptors, LMCs express voltage-dependent K^+^ conductances [15] but their membrane shows little or no rectification to injected current [39]. Consequently, the input resistance is likely to be a good estimate of the LMC membrane resistance, implying that previous calculations of energy consumption and the energy efficiency of information coding [31] are likely to be accurate.

### 4.3 Robustness and limitations

Our conclusions do not depend critically upon the values we used to parameterise our model of the blowfly photoreceptor membrane. Increasing or decreasing total conductance, number of gating particles, activation time constant, *a* and *b* parameters in Table 1, and K^+^ leak conductance by up to 20%, and adjusting the leak conductance to obtain a dark resting potential of −60 mV, produces membrane resistances that are always 1.9-2.5 higher than input resistances. The corresponding membrane bandwidths are 2.4- to 3.4-fold higher than each corresponding passive membrane. When depolarised to −37 mV by increasing the light-induced conductance, the bandwidths are 1.35- to 1.8-fold higher than each corresponding passive membrane. Thus, changes in experimental measures of the biophysical parameters governing the behaviour of the voltage-dependent conductances are unlikely to alter our conclusions.

Incorporating the additional molecular processes involved in phototransduction into our photoreceptor energy consumption calculation does not alter it substantially. At the highest light levels, the penultimate stage of phototransduction, PIP_2_ signalling [40], consumes 4.2 × 10^8^ ATP/s, equivalent to *∼*112-140 ATP molecules per transduced photon (see Supplement 1), which is less than 5% of the cost of maintaining ionic gradients. Thus, a fly’s rhabdomeric photoreceptor is similar to a mouse’s ciliary rod photoreceptor in which less than 10% of metabolic energy is consumed by the biochemical processes of phototransduction under most conditions; the exception being when bright light closes all of a rod’s light-gated channels to save energy during the day [41, 42]. These findings emphasize that ion flux across the membrane is the primary energy consuming process in a neuron [31, 1, 29, 8].

Inclusion of the Na^+^/Ca^2+^ exchanger, which exchanges 1 Ca^2+^ for 3 Na^+^ ions in microvilli [43, 44], would produce an additional inward current. Assuming that *∼*26% of the LiC in blowfly photoreceptors is Ca^2+^ mediated, as in fruit fly photoreceptors [45], this current would be *∼*13% of the LiC, producing a small increase in impedance without affecting the cost. Likewise, incorporating the conductances in the photoreceptor terminal [30, 46] or electrical coupling between photoreceptor axons [47] is unlikely to have a substantial effect on energy consumption.

### 4.4 Functional significance of tuning membrane bandwidth

Expressing two DRs with different time constants and voltage operating ranges has clear advantages for blowfly photoreceptors. The FDR activates close to the resting potential, and improves bandwidth at all light intensities by generating negative feedback that reduces impedance at frequencies below *∼*100 Hz and by shunt peaking [21]. The SDR activates at more depolarised potentials, and its negative feedback lowers impedance below *∼*10 Hz at high light intensities, thereby producing most of the low frequency roll-off of the membrane frequency response. The progressive shift from low-pass filtering at low light intensities when the signal-to-noise ratio (SNR) is low, to band-pass filtering at high light intensities when the SNR is high, is optimum for coding visual stimuli with natural image statistics within a limited dynamic range [35, 36]. Thus, by activating progressively with depolarisation the FDR and SDR contribute to the energy-efficient coding of natural images across a broad range of light intensities. Importantly, membrane filtering retains two key properties of filtering by a passive membrane. One is stability, the membrane produces bounded responses to bounded current injections, and the other is minimum phase, the group delay is minimum for filters with the same impedance gain [21].

Although the membrane bandwidth of blowfly R1-6 photoreceptors is 3- to 4-fold greater than the bandwidth of the LiC [24, 48], several lines of evidence demonstrate that a greater membrane bandwidth important. (i) Investing in a high bandwidth membrane is expensive but fly photoreceptors do precisely that when additional bandwidth is needed [31, 29]. (ii) Across different fly species, photoreceptor membrane dynamics are tuned to the bandwidth of LiC so that those operating with high SNR and information rates invest in expensive wide-bandwidth membranes [49, 29]. (iii) In blowfly retina, the responses of frontal R1-6 photoreceptors are faster than those at the back or the side. This gradient in voltage response bandwidth is matched by a gradient in membrane bandwidth, with frontal photoreceptors having both the fastest voltage responses and the fastest membranes, and those at the side having the slowest voltage responses and the slowest membranes [48]. (iv) In male houseflies, the love-spot (a region in the eye specialised to detect mates during chasing) the voltage light responses of male R1-6 photoreceptors have higher bandwidth than the equivalent photoreceptors in the female. This is due both to improvements in LiC bandwidth and membrane bandwidth [50]. (v) Finally, during light adaptation, most fly photoreceptors—including R1-6 in the blowfly — improve both LiC bandwidth and membrane bandwidth. In blowfly R1-6 photoreceptors, membrane bandwidth changes with adaptation to maintain a constant SNR at its cut-off frequency [24].

Not all insect photoreceptors contain voltage-dependent ion channels with properties similar to the DRs that blowfly and other fast flying flies use to regulate membrane gain and bandwidth. Indeed, photoreceptors from the retinas of different species, or even from within the same retina [33, 51], express different types of voltage-dependent ion channels. Slow moving/nocturnal species having slower light responses than those from diurnal, fast moving species [49, 52, 53], and their photoreceptors likely save energy by expressing inactivating conductances and having low bandwidth [49, 54, 18]. In such photoreceptors, inactivating conductances may play the same role as DRs in blowflies; producing negative feedback and shunt peaking, while preventing a large increase in membrane resistance to save energy.

### 4.5 Implications for non-spiking neurons and dendrites

The voltage-dependent K^+^ conductances of other neurons or compartments within neurons may also be increasing the bandwidth of analogue signals in an energy aware way. For example, rods and cones express hyperpolarization-activated cyclic nucleotide-gated (HCN) channels that open with increasing light intensity. These channels reduce impedance and improve the frequency response by acting as an inductance [55]. Inner hair cells of the mammalian cochlea also express DRs that decrease the time constant of their response to injected current [56]. Dendrites are known to express different types of voltage-dependent conductances [11], and their effect on filtering has been widely studied with experimental and computational techniques [57, 58, 59]. However, the implications for energy efficiency have not been considered in these other neurons and neuronal compartments. Our modelling of the action of voltage-dependent K^+^ conductances in blowfly photoreceptor membrane suggests that, in addition to regulating membrane gain and frequency response, they could increase the energy efficiency of analogue signalling.

## Data accesibility

The code underlying the findings described in the manuscript (the Python package pHHotoreceptor and scripts generating figures) can be downloaded from *https://github.com/fjhheras/phhotoreceptor*

## Acknowledgements

The authors thank R.C. Hardie, M. Lengyel, K.D. Longden, D.G. Stavenga and M. Wicklein for discussions and feedback on previous versions of this manuscript.

## Authors’ contributions

All authors conceived the study. J.A. performed experiments and fitted conductance properties, F.J.H.H wrote code and conducted analyses. F.J.H.H, S.B.L and J.E.N. wrote the manuscript with input from J.A. All authors gave final approval for publication.

